# Immune-Epithelial Dynamics and Tissue Remodeling in Chronically Inflamed Nasal Epithelium via Multi-scaled Transcriptomics

**DOI:** 10.1101/2023.07.01.547352

**Authors:** Guanrui Liao, Tsuguhisa Nakayama, Ivan T. Lee, Bokai Zhu, Dawn. T. Bravo, Jonathan B. Overdevest, Carol H. Yan, David Zarabanda, Philip A. Gall, Sachi S. Dholakia, Nicole A. Borchard, Angela Yang, Dayoung Kim, Zara M. Patel, Peter H. Hwang, Dhananjay Wagh, John Coller, Katie Phillips, Michael T. Chang, Matt Lechner, Qin Ma, Zihai Li, Garry Nolan, Dan H. Barouch, Jayakar V. Nayak, Sizun Jiang

## Abstract

Chronic rhinosinusitis (CRS) is a common inflammatory disease of the sinonasal cavity that affects millions of individuals worldwide. The complex pathophysiology of CRS remains poorly understood, with emerging evidence implicating the orchestration between diverse immune and epithelial cell types in disease progression. We applied single-cell RNA sequencing (scRNA-seq) and spatial transcriptomics to both dissociated and intact, freshly isolated sinonasal human tissues to investigate the cellular and molecular heterogeneity of CRS with and without nasal polyp formation compared to non-CRS control samples. Our findings reveal a mechanism for macrophage-eosinophil recruitment into the nasal mucosa, systematic dysregulation of CD4+ and CD8+ T cells, and enrichment of mast cell populations to the upper airway tissues with intricate interactions between mast cells and CD4 T cells. Additionally, we identify immune-epithelial interactions and dysregulation, particularly involving understudied basal progenitor cells and Tuft chemosensory cells. We further describe a distinct basal cell differential trajectory in CRS patients with nasal polyps (NP), and link it to NP formation through immune-epithelial remodeling. By harnessing stringent patient tissue selection and advanced technologies, our study unveils novel aspects of CRS pathophysiology, and sheds light onto both intricate immune and epithelial cell interactions within the disrupted CRS tissue microenvironment and promising targets for therapeutic intervention. These findings expand upon existing knowledge of nasal inflammation and provide a comprehensive resource towards understanding the cellular and molecular mechanisms underlying this uniquely complex disease entity, and beyond.

## Introduction

Chronic rhinosinusitis (CRS) is a recondite and heterogeneous inflammatory disease of the nasal and sinonasal cavities. Epidemiologic studies estimate the global prevalence of CRS to be approximately 12% (1, 2) with patient-rated symptom severity akin to heart disease and chronic back pain (1). CRS can be classified into two major subtypes based on the presence or absence of nasal polyps: CRS with nasal polyps (CRSwNP) and CRS without nasal polyps (CRSsNP). Of the total CRS population, CRSsNP typically accounts for 75-80% of patients seen vs. 20-25% for CRSwNP (3), although this proportion varies regionally However, CRSwNP in particular is associated with higher disease burden from obstructive, eosinophil-rich, nasal polyposis and sinonasal outflow tract inflammation and infection, leading to an increased likelihood of recalcitrant symptoms such as sinus headaches, olfactory loss, and recurrent sinusitis. The pathogenesis of CRSwNP involves both innate and acquired Th2-immunity mediated by the nasal epithelium/mucosa due to stimulation by extrinsic antigens, but the interaction between immune cells, epithelial cells, and key molecular determinants driving disease progression, remains elusive.

The dynamic crosstalk between immune-epithelial systems plays a critical role in the pathogenesis of many diseases, including CRS (4–6). In addition to its role as a physical barrier against environmental challenges from pathogens, airborne particulates and allergens, the nasal epithelium generates cell-derived cytokines and chemokines involved in mediating autocrine and paracrine signaling. These events lead to recruitment of diverse myeloid and lymphoid immune cells, that in turn release molecular mediators that invigorate or blunt downstream epithelial and immune cell functions, thus orchestrating signature acute vs. chronic inflammation. This subtle interplay between epithelial and immune cells is often bidirectional within the native tissue microenvironment, and involves multiple participants.

T cells naturally play a crucial role in the adaptive immune response, and are central for regulation of the immune-epithelial interactions responsible for CRS pathogenesis. In particular, CD4+ T cells can differentiate into various subpopulations based on the cytokine environment encountered. CD4+ Th2 cells produce cytokines such as interleukin-4 (IL-4), IL-5, and IL-13, that recruit and activate eosinophils and mast cells which have been well-established to play significant roles in the pathophysiology of CRSwNP (7). CD8+ T cells eliminate infected or damaged cells, with their specific contributions to CRS less appreciated.

Mast cells, another key player in the pathogenesis of CRS, are involved in innate immunity release of a range of inflammatory mediators, including histamine, prostaglandins, and leukotrienes (8). Elevated mast cell number in CRSwNP has been reported, with their activation linked to the presence of cytokines and chemokines that promote eosinophilic inflammation (9).

Basal cell differentiation is an important factor in the pathogenesis of CRS. The sinonasal epithelium is comprised of several distinct cell types, including basal cells along the epithelial basement membrane, as well as differentiated ciliated cells, and goblet cells oriented towards the airway lumen. Basal cell hyperplasia, a rise in basal cell numbers through cell division, has been detected in patients with CRSwNP (10, 11), although the physiological relevance and consequence has been unclear. Basal cells differentiate into the other major ciliated and goblet/secretory epithelial cell types in response to environmental stressors (12, 13), but whether this process in fostering the development of CRS through priming of epithelial-immune exchange is entirely uncertain. We have previously described prominent type II responses in macrophages, and laid the groundwork to better assess distinctive inflammatory and epithelial cells and their contributions to type II inflammatory profiles in CRSwNP patients (14).

A better understanding of these mechanisms in situ is crucial for the development of more targeted and effective treatments for this common, challenging and debilitating upper airway disease. To achieve this, we applied single-cell sequencing to uncover the phenotypic composition and functional aspects of a discovery CRS clinical cohort (Fig.1A), and orthogonally utilized spatial transcriptomics to interrogate a validation CRS cohort (Fig.1A) to untangle the key players and epithelial-immune interactions within inflamed nasal tissues, including CRSwNP. We envision such a resource will also be broadly applicable to the multitude of other nasal inflammatory diseases.

**Figure 1:**
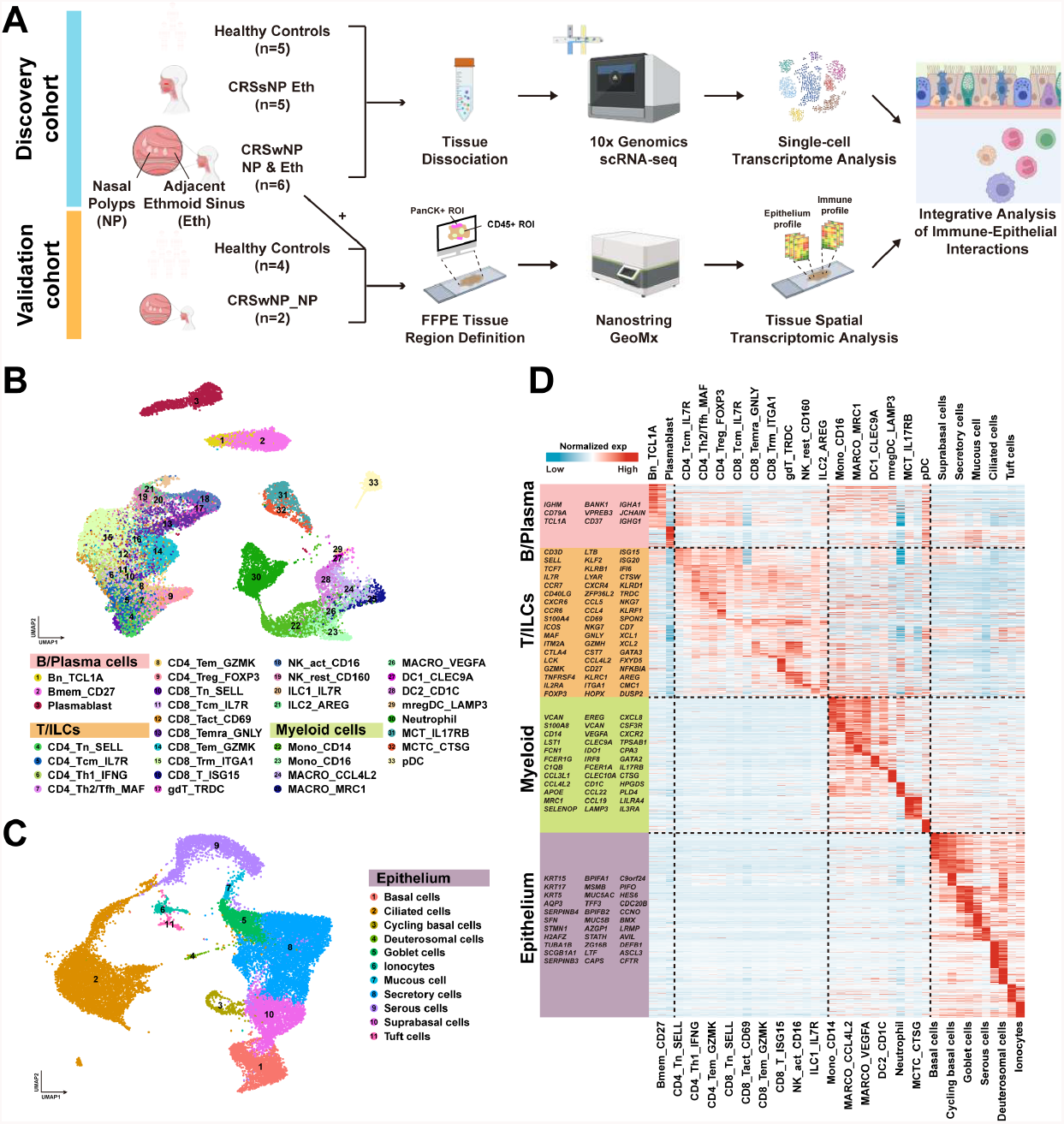
Comprehensive Single-Cell Transcriptomic Analysis Reveals the Complex Immune and Epithelial Microenvironment in CRS. (A) Schematic representation of the experimental workflow for the analyses conducted on CRS and control samples in the discovery and validation cohorts. CRSsNP - CRS without nasal polyp; CRSwNP - CRS with nasal polyps;(B) Uniform Manifold Approximation and Projection (UMAP) plot depicting 3 major cell types and 33 subtypes within the immune microenvironment of CRS, color-coded by cell type. (C) UMAP plot depicting the 11 epithelial cell types identified. (D) Heatmap depiction of the expression patterns of signature genes across the immune and epithelial cell types identified in panels (B) & (C), respectively.

## Results

### Single-Cell Transcriptomic Analysis of the CRS Microenvironment

We utilized single-cell transcriptomics for an in-depth analysis of the CRS epithelial and immune landscape on an initial discovery cohort of rigorously-selected patients (n = 5 healthy controls, n = 5 CRSsNP, n = 6 CRSwNP for both the NP and adjacent non-polyp ethmoid sinus mucosa, see Methods) (Fig.1A and S1A). We first identified the major immune cell types within the upper airway microenvironment (Fig.1B), as B, T, and myeloid lineages. The origins of the 32,775 total cells were displayed in a UMAP plot, with tissue types and patient samples color-coded (Fig.S1B) as well as representative genes across the immune cell repertoire(Fig.S1C). We further resolved 11 cell types (21,833 cells in total) present within the upper airway human tissue samples across healthy and CRS samples, including secretory, ciliated, basal, goblet, tuft and other epithelial cell types (Fig.1C). The epithelial cell origins were presented in a separate UMAP plot, with tissue types and patient samples colorcoded (Fig.S1D), and representative canonical marker genes across the epithelial cell repertoire depicted (Fig.S1E) Signature gene expression patterns were further discriminated across both immune and epithelial cell types (Fig.1D), to gain detailed insight into the complex cellular composition and states in CRS tissues.

### Macrophage Polarization in CRS Nasal Polyps

Given the postulated role of myeloid cells in CRS (10, 14), we further stratified the myeloid cluster into subtypes, including macrophages, monocytes, and dendritic cells (DCs) (Fig.2A). These subtypes were well represented across the healthy and CRS samples (Fig.S2A). We quantified the percent composition of the three main subtypes of macrophages identified (CCL4L2, MRC1, VEGFA), and observed little change between numbers of the more M1-like macrophages state (Fig.2B, left panel), while macrophage subtypes polarized towards M2-like gene expression were consistently and significantly elevated in CRSwNP compared to healthy controls or CRSsNP (Fig.2B, middle and right panels). A similar analysis was performed for the other myeloid cells without any notable differences (Fig.S2B). These results suggested that macrophage cell states, and not merely quantities, are dysregulated in CRSwNP. We thus performed differential gene analysis to identify differentially expressed genes (DEGs) responsible for the cell state differences between the macrophages from CRSsNP and CRSwNP tissues (Fig.2C). Amongst them were genes associated with antigen presentation, complement pathway activation, and chemokines linked to immune cell recruitment and activation (Fig.2C and Fig.S2C). Scoring of immunosuppressive M2 activity through a pre-curated set of genes (15, 16) confirmed the increased frequency of M2-polarized in polyp tissue from single-cell RNA-seq (Fig.2D) and spatial (Fig.S2D), compared to non-polyp ethmoid tissue.

**Figure 2:**
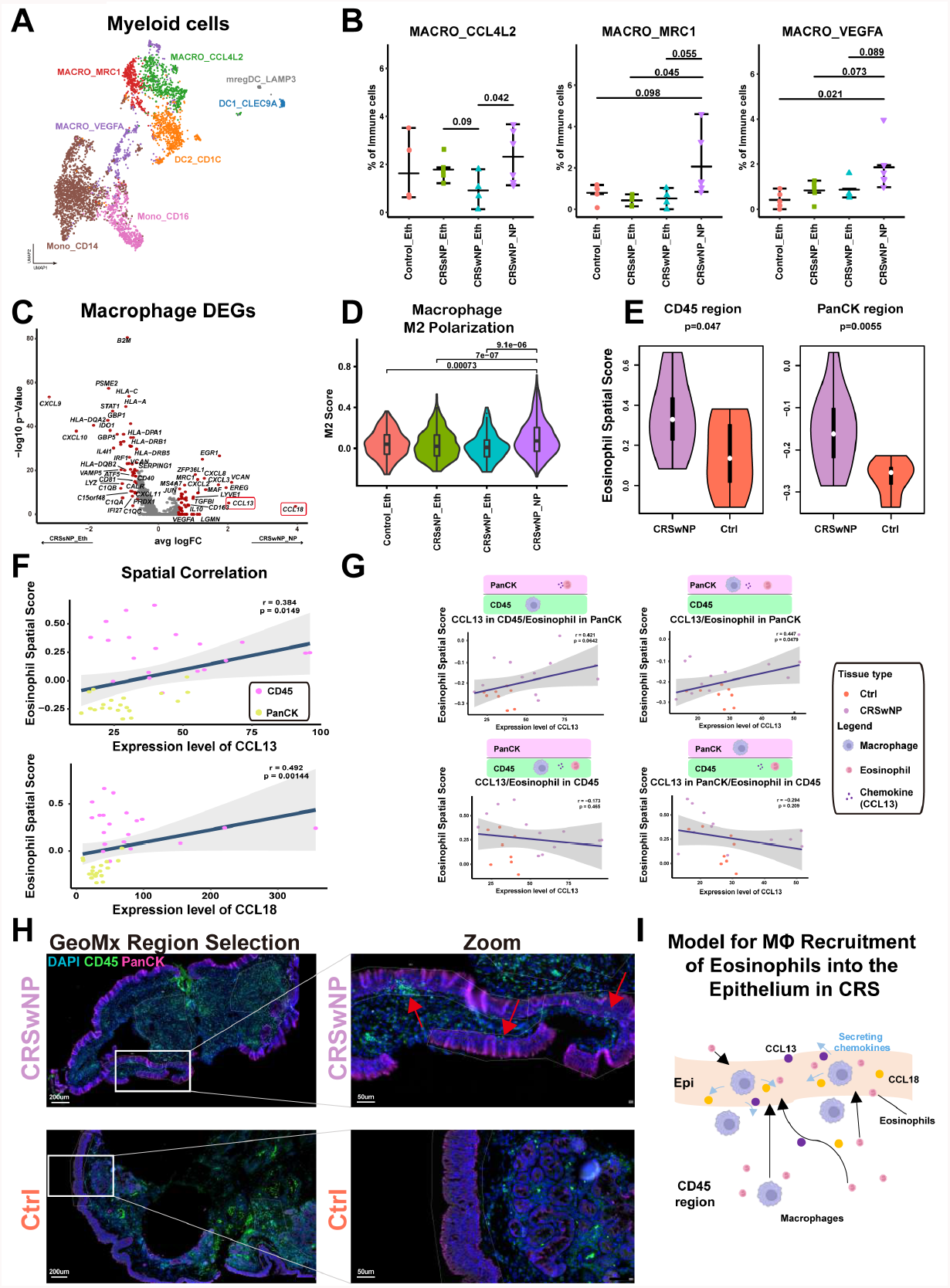
Polarization of Macrophages to M2 Phenotype Drives Type 2 Inflammation in CRS Nasal Polyps. (A) UMAP plot depicting subtypes and corresponding annotations of myeloid cells in CRS and healthy control samples. (B) Comparison of macrophage cell fractions between CRS and control samples using the Wilcoxon test (two-sided). (C) Volcano plot displaying differentially expressed genes in macrophages between CRSsNP and CRSwNP, with the most significant genes indicated in red (|*Foldchange*| > 1.5), including CCL13 and CCL18. (D) Violin plots illustrating M2 scores for macrophages across CRS and control samples, with comparisons performed using the Wilcoxon test (two-sided) and p values indicated. (E) Violin plots comparing eosinophil spatial signature expression scores between CRS nasal polyps (purple) and healthy control samples (orange) in spatial transcriptomics GeoMx data within CD45+ regions (left panel) and PanCK+ regions (right panel), with comparisons performed using the Wilcoxon test (two-sided) and p values indicated. (F) Scatter plots demonstrating the correlation between *CCL13* (upper panel) or *CCL18* (lower panel) mRNA expression levels in situ, and eosinophil spatial signature expression scores in GeoMx data, with data origins colored to indicate CD45+ regions (magenta) and PanCK+ regions (yellow). The data was fitted using a linear regression model, with blue lines indicating the mean and grey regions highlighting the 95% confidence intervals. The regression index and p values are provided within the plots. (G) Scatter plots illustrating the correlation between *CCL13* expression levels and eosinophil signature scores in CD45+ or PanCK+ regions of GeoMx Spatial Transcriptomics acquisition, with sample origins color-coded to represent CRS nasal polyps (purple) and healthy control samples (orange). Diagrams above the scatter plots indicate regions where *CCL13* and eosinophil spatial gene signatures were captured. (H) Representative multiplexed immunofluorescence images from the GeoMX spatial transcriptome acquisition from a CRSwNP sample (upper panel) and a healthy control sample (lower panel). Red arrows highlight immune infiltration into the epithelial region, as indicated by CD45-positive cells within the PanCK region. White outlines indicate the region from which the transcriptome was extracted from for the GeoMx experiment. (I) The proposed model in which macrophages secreting *CCL13*/*CCL18* chemokines attract eosinophils to infiltrate the epithelium in CRS nasal polyps.

### Macrophage Recruitment of Eosinophils in CRS Through *CCL13* and *CCL18*

Given the known role of CCL13 and CCL18 in CRSwNP (Fig.2A) for the recruitment of monocytes, including eosinophils (17, 18), we first confirmed that eosinophils were increased in nasal polyp tissue compared to control ethmoid tissues via spatial transcriptomics (Fig.2E). This leverages upon the intact tissue microenvironment preserved by spatial transcriptomics, since single-cell dissociation approaches can often result in the loss of specific cell-types (19). We next tested the hypothesis that CCL13 and CCL18 were involved in the recruitment of eosinophils by macrophages (14). From our spatial transcriptomics data, we observed significant correlations in the expression of both chemokines with heightened eosinophilic signatures in both the immune and epithelial tissue regions (Fig.2F). We next postulated that a location-based pairwise spatial analysis of these signatures would enable insights into the dynamics of eosinophil recruitment by macrophages. We observed a strong correlation between *CCL13* and *CCL18* expression with the influx of eosinophils in the pan-cytokeratin (PanCK)-positive epithelial, but not CD45-positive immune regions (Fig.2G and Fig.S2E). Similarly, the correlative expression of *CCL18* and its receptor, *CCR2*, also supported a case of directionality in the attraction of eosinophils into the epithelial region, but not the immune region, of the CRS nasal tissues.

Representative immunofluorescence images from the tissues stained for GeoMx, and regions defined for spatial transcriptomics data collection, further substantiated the localization of immune cell infiltration into the epithelial regions in CRSwNP, but not healthy control mucosal tissues (Fig.2H). Taken together, these results suggest a model, in which macrophages secreting *CCL13*/*CCL18* in CRSwNP are directing recruitment to, and subsequent trafficking of, eosinophils into the nasal epithelium in CRSwNP disease (Fig.2I) (14).

### Immunosuppressive CD4+ and CD8+ T Cell Responses Predominate in Nasal Polyps

Detailed analysis of CD4+ T cells and their subtypes revealed several categories represented across the control and CRS samples (Fig.3A and Fig.S3A). We identified an enrichment of CD4+ T effector memory (TEM), Th2, and T regulatory (Treg) CD4+ subtypes in CRSwNPs, and a depletion of Th1 CD4+ cells as previously described (Fig.3B and Fig.S3B) (20). Differential gene expression analysis and pathway enrichment analysis demonstrated significant differences between CD4+ T cells within the CRSsNP and CRSwNP microenvironment (Fig.3C and Fig.3D), especially when compared against healthy controls (Fig.S3C). We confirmed the increased CD4+ T cell immunosuppression within CRSwNP compared to CRSsNP as demonstrated by Th2-skewed inflammation from the scRNAseq cohort (Fig.3E), a reduction of immune cells related to the Th1 pathway, and an increase of immune cells towards the Th2 pathway from spatial transcriptomics (Fig.S3D).

**Figure 3:**
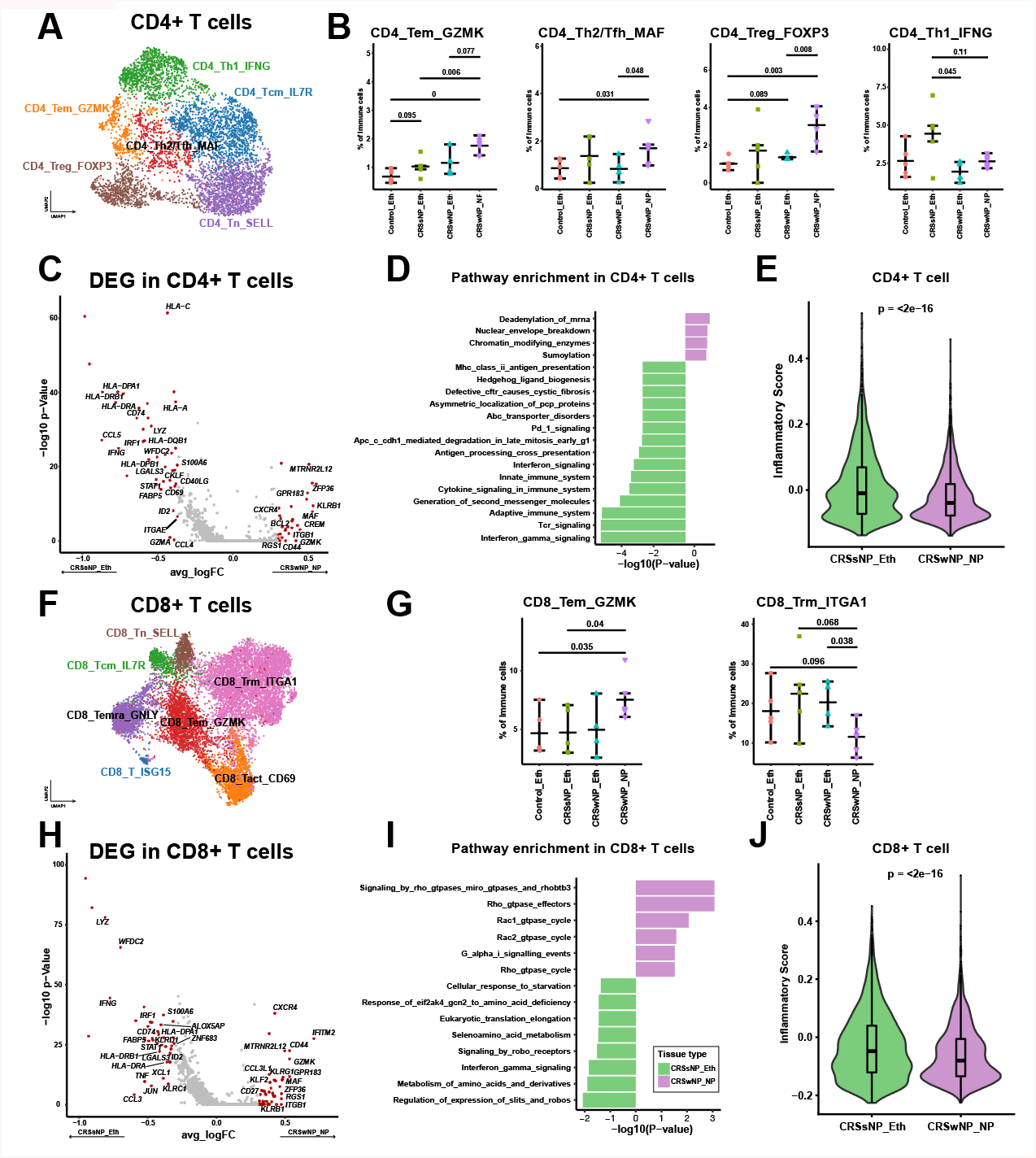
Regulatory CD4+ and CD8+ T Cells Predominate in Nasal Polyps. (A) UMAP plot illustrating subtypes and corresponding annotations of CD4+ T cells in CRS and control samples. (B) Comparison of CD4+ T cell fractions between CRS and control samples using the Wilcoxon test. (C) Volcano plot displaying differentially expressed genes in CD4+ T cells between CRSsNP and CRSwNP, with the most significant genes indicated in red (|*F oldchange*| > 1.25). (D) Pathways enriched in CD4+ T cells from CRSsNP and CRSwNP, based on GSEA analysis using the REACTOME gene set. (E) Violin plots illustrating CD4+ T cell inflammatory signature expression scores in CRS and control samples, with comparisons performed using the Wilcoxon test (two-sided) and p values indicated. (F) UMAP plot depicting subtypes and corresponding annotations of CD8+ T cells in CRS and control samples. (G) Comparison of CD8+ T cell fractions between CRS and control samples using the Wilcoxon test (two-sided). (H) Volcano plot showing differentially expressed genes in CD8+ T cells between CRSsNP and CRSwNP, with the most significant genes indicated in red (|*F oldchange*| > 1.25). (I) Pathways enriched in CD8+ T cells from CRS nasal polyps versus CRS without nasal polyps, based on GSEA analysis using the REACTOME gene set. (J) Violin plots presenting CD8+ T cell inflammatory signature expression scores in CRS and control samples, with comparisons performed using the Wilcoxon test (two-sided) and p values indicated.

Similarly, we investigated and identified lymphocyte subtypes and corresponding annotations in CD8+ T cells in both CRS and control samples (Fig.3F and Fig.S3E). Similar to our CD4+ T cell analysis, we also identified the enrichment of TEMs in the CRSwNP samples, along with a reduction in CD8+ resident memory T cell phenotypes (Fig.3G and Fig.S3F). Differential gene expression analysis and pathway enrichment analysis also discriminated significant differences in CD8+ T cells between the CRSsNP and CRSwNP microenvironment (Fig.3H and Fig.3I), along with altered inflammation (Fig.3J), in line with the CD4+ T cell findings (Fig.3E and Fig.S3D). These results support a model in which suppressor and regulatory T cells, including players involved in a type II immune response and Tregs, are responsible for the unique chronic inflammatory features of CRSwNP compared to CRSsNP (21).

### Mast Cell Enrichment and Type 2 Immune Responses in Nasal Polyps

Given the intricate relationship between mast cells (MCs) and the type II immune response in T cells, we sought to better define the possible role of mast cells in CRSwNP disease (9). We observed two major subtypes of mast cells, stratified into 1) epithelial MCs expressing TPSAB1 tryptase without CMA1 chymase, with high expression of interleukin 17 Receptor B (termed MCT_IL17RB), and 2) subepithelial MCs with high expression of the tryptase protease, along with Cathepsin G (CTSG) and chymase (termed MCTC_CTSG) (Fig.4A). Both MC subtypes were found to be enriched in CRSwNP compared with other sample types (Fig.4B). The expression patterns of signature genes in these two mast cell subtypes were visibly distinct (Fig.4C), channeling the nuanced different cell states and functions within the CRSwNP tissue microenvironment. We therefore postulated that these mast cells subtypes may have distinct roles in the recruitment and interaction with key immunocyte players within the CRSwNP tissue microenvironment. We tested this hypothesis via Ligand-Receptor (L-R) analysis and identified several pathways for immune and tissue remodeling related to CD4+ T cells, including *IL2, OX40, CCL, EPHB, PROS, IL4*/*IL13, PARs, CD22, ICAM, SEMA7, LIFR, CLEC*, and *OSM* (Fig.4D). Of particular interest were the key cytokine mediators in Type II inflammation: *IL4* and *IL13* (Fig.4D), which was predominantly expressed by MCs in our study (Fig.4E), and were implicated in MC and CD4+ T cell interactions in CRSwNP and not CRSsNP (Fig.4F). The CSF2 signaling pathway served as a control (Fig.4F). While similar trends were observed in each of the mast cell clusters (Fig.S4A-E), the MCT_IL17RB mast cells exhibited a higher potential for immune interaction in CRSwNP as previously reported (22).

**Figure 4:**
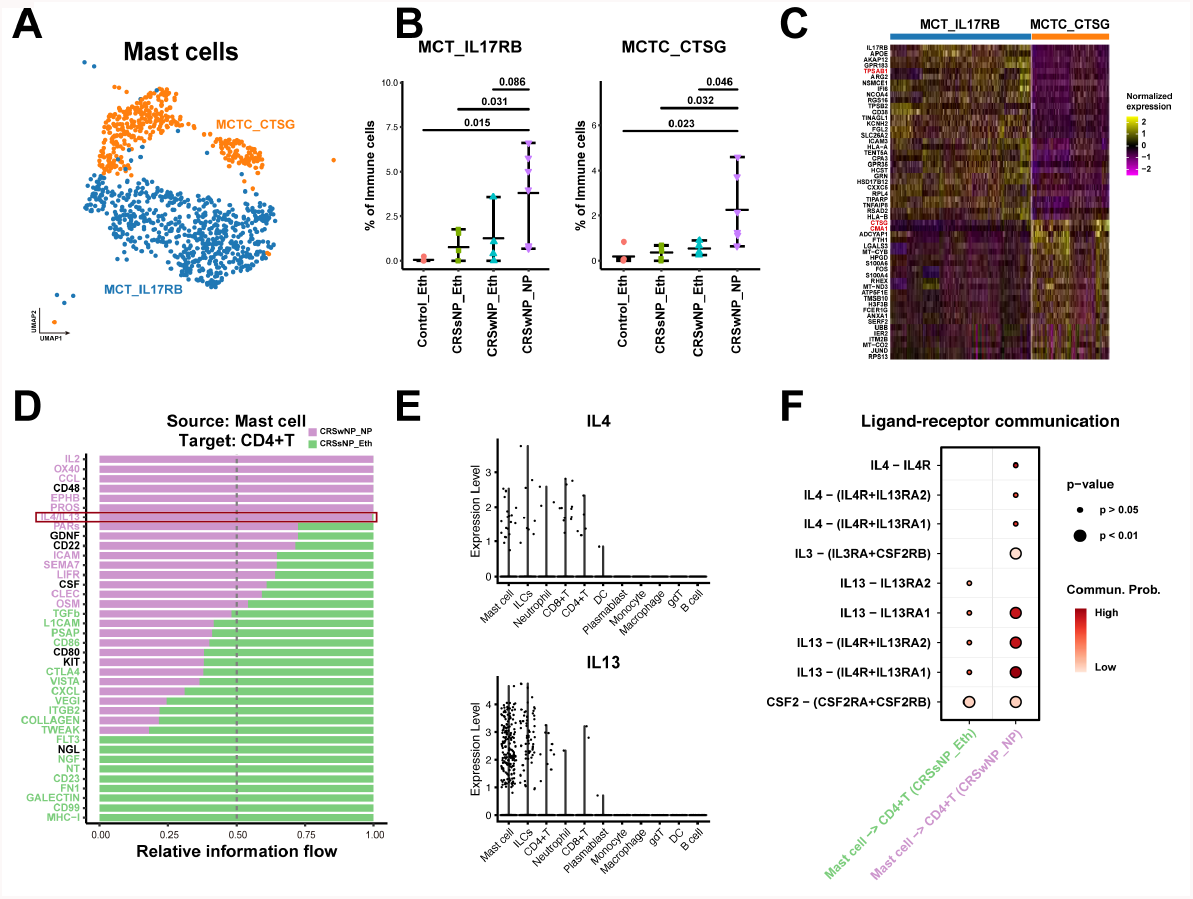
Mast Cell Enrichment in Nasal Polyps Correlates with Type 2 Immune Responses. (A) UMAP plot illustrating subtypes and corresponding annotations of mast cells in CRS and control samples. (B) Comparison of mast cell subtype fractions between CRS and control samples using the Wilcoxon test (two-sided). (C) Heatmap displaying normalized expression level of signature genes in the mast cell subtypes identified. (D) Ligand-receptor (L-R) interactions identified between mast cells and CD4+ T cells in CRSwNP (purple) and CRSsNP (green). L-R pairs with purple bars crossing the 0.5 dotted line indicate predominance in CRSwNP, while those with green bars crossing the dotted line indicate predominance in CRSsNP. Significant interactions are color-coded accordingly (p < 0.05, Wilcoxon test, two sided). (E) Scatter plots depicting IL4 and IL13 expression levels in various immune cells, and their dominant expression in Mast cells. (F) Dot plot demonstrating the significance and strength of IL4/IL13-related ligand-receptor interactions between mast cells and CD4+ T cells in CRSwNP (purple) and CRSsNP (green).

### Identification of Key Players in the Immune-Epithelial Crosstalk and Remodeling in CRSwNP

Given the data from our work and others on the emerging evidence of immune-epithelial crosstalk and remodeling in multiple diseases (5, 6), including CRSwNP (4), we postulated that quantifying cell abundance correlations between immune and epithelial cell subsets in CRS and control samples would identify potential key players in this axis. Our analysis revealed a key cluster of epithelial and immune cell types that were strongly correlated with each other, indicative of their potential interplay in the epithelial-immune crosstalk and remodeling in CRSwNP (Fig.5A; black box). We specifically observed the enrichment of Tuft cells, cycling basal cells, and suprabasal cells as enriched in CRSwNP polyps, and the conversely depletion of FoxJ1 low ciliated cells, mucous cells and serous cells in CRSwNP polyps (Fig.5B and Fig.S5). These results warrant further investigation of Tuft cells and basal cells as key players in mediating the immune-epithelial crosstalk and attraction of immune infiltrates in the context of chronic inflammation with nasal polyps formation.

**Figure 5:**
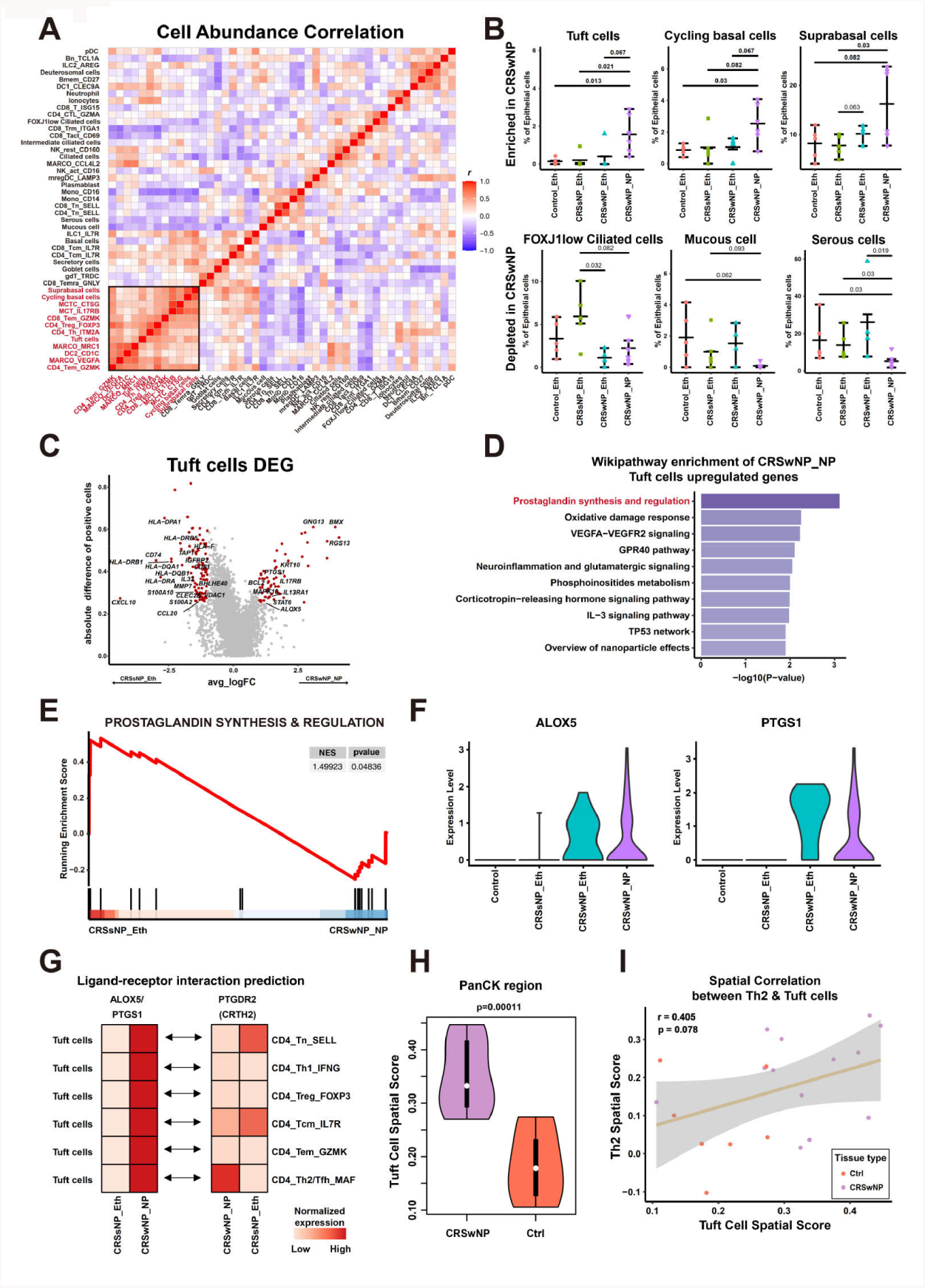
Tuft Cells in Nasal Polyps Correlate with Th2 Cells. (A) Heatmap illustrating cell abundance correlations between immune and epithelial cell fractions. (B) Comparison of epithelial cell subtype fractions between CRS and control samples using the Wilcoxon test (two-sided). (C) Volcano plot depicting differentially expressed genes in Tuft cells between CRSwNP and CRSsNP. The most significant genes are highlighted in red (|*F oldchange*| > 2 and Δpct > 0.25). (D) Pathways enriched in Tuft cells from CRSwNP and CRSsNP, based on WIKIPATHWAY enrichment analysis. (E) Enrichment plot of the prostaglandin synthesis and regulation pathway in Tuft cells from CRSwNP versus CRSsNP, using GSEA analysis with the WIKIPATHWAY gene set. The enrichment score and p-value are indicated in the plot. (F) Violin plot displaying expression levels of ALOX5 and PTGS1 in CRS and control samples. (G) Heatmap presenting mean expression levels of ALOX5/PTGS1 ligands in Tuft cells and mean expression levels of their PTGDR2 receptor in CD4+ T cell subsets in CRSwNP and CRSsNP. (H) Violin plots comparing Tuft cell spatial gene signature expression scores between CRSwNP (purple) and healthy controls (orange) in GeoMx spatial transcriptomics data within PanCK+ tissue regions. (I) Scatter plot and regression line illustrating the correlation between Tuft cell spatial gene signature expression scores in PanCK+ regions and Th2 cell spatial gene signature expression scores in CD45+ regions. Dots are colored to represent patient sample origins. The grey region indicates the confidence interval. The regression index and p-values are shown in the plots.

We identified multiple cell-signaling pathways (including G protein, Tyrosine Kinase, and MAP Kinase members), anti-apoptotic genes (i.e. *BCL2*), and cytokine pathways (i.e. *IL17RB, IL13TA1, STAT6*) upregulated in CR-SwNP (Fig.5C). Conversely, components of the antigen-presentation pathway were upregulated in CRSsNP (Fig.5C), implicating different cell states of the tuft cells in CRSwNP as opposed to CRSsNP. We next identified additional pathways enriched in Tuft cells in CRSwNP, particularly the prostaglandin pathway (Fig.5D), an inflammatory pathway previously not described in the context of CRS. Gene Set Enrichment Analysis orthogonally confirmed the activation of the prostaglandin pathway in CRSwNP (Fig.5E), along with the expression of key members of this pathway, ALOX5 and PTGS1, in CRSwNP polyps and adjacent ethmoid tissues, suggestive of high prostaglandin pathway activity in Tuft cells within and outside of nasal polyps (Fig.5F). Ligand-receptor analysis revealed significant pairing of tuft cell interactions with Th2 CD4+ T cell recruitment in CRSwNP, as well as depletion of naive and central memory CD4+ T cells (Fig.5G), in line with our abundance correlative analysis (Fig.5A). We next confirmed the increased density of tuft cells within the CRSwNP epithelial layer in situ through spatial transcriptomics (Fig.5H), to support the hypothesized tissue interactions between Tuft cells in the PanCK+ region and Th2 CD4+ T cells in CD45+ region of the CRSwNP tissue (Fig.5I). These results strongly implicate chemosensory tuft cells as one of the epithelial mediators of immune cell recruitment, including recruiting CD4+ Th2 cells into the CRSwNP inflammatory microenvironment to prime Type II inflammation.

### Identification of a Basal Cell Trajectory That Drives Key Epithelial-Immunologic Remodeling for Nasal Polyp Formation

We finally investigated the role of basal cells, which were also implicated as critical in CRSwNP epithelial-immune remodeling (Fig.5A and 5B). We observed differences in the expression of key genes between suprabasal cells and cycling basal cells (Fig.6A and Fig.S6A), which included a sizable overlap of key genes upregulated in CRSwNP compared to CRSsNP (Fig.S6B). Given the proliferative and developmental potential of basal cells, including towards differentiated and/or specialized cell fates, we postulated that a cell trajectory analysis would allow us to track differentiation states of the basal cells. Using the pseudotime analysis, we confirmed that undifferentiated basal cells tend to be present at a much earlier pseudotime point, followed by a bifurcation in basal cell developmental trajectory, which we termed Cell-fate1 and Cell-fate2 (Fig.6B). We observed an enrichment of basal cells from CRSwNP patients in Cell-fate2, while those from control and CRSsNP tissues were associated with Cell-fate1 (Fig.6C-D), suggesting disparate outcomes and cell states for the differentiated basal cells in CRSwNP upper airway milieu compared to the CRSsNP microenvironment.

**Figure 6:**
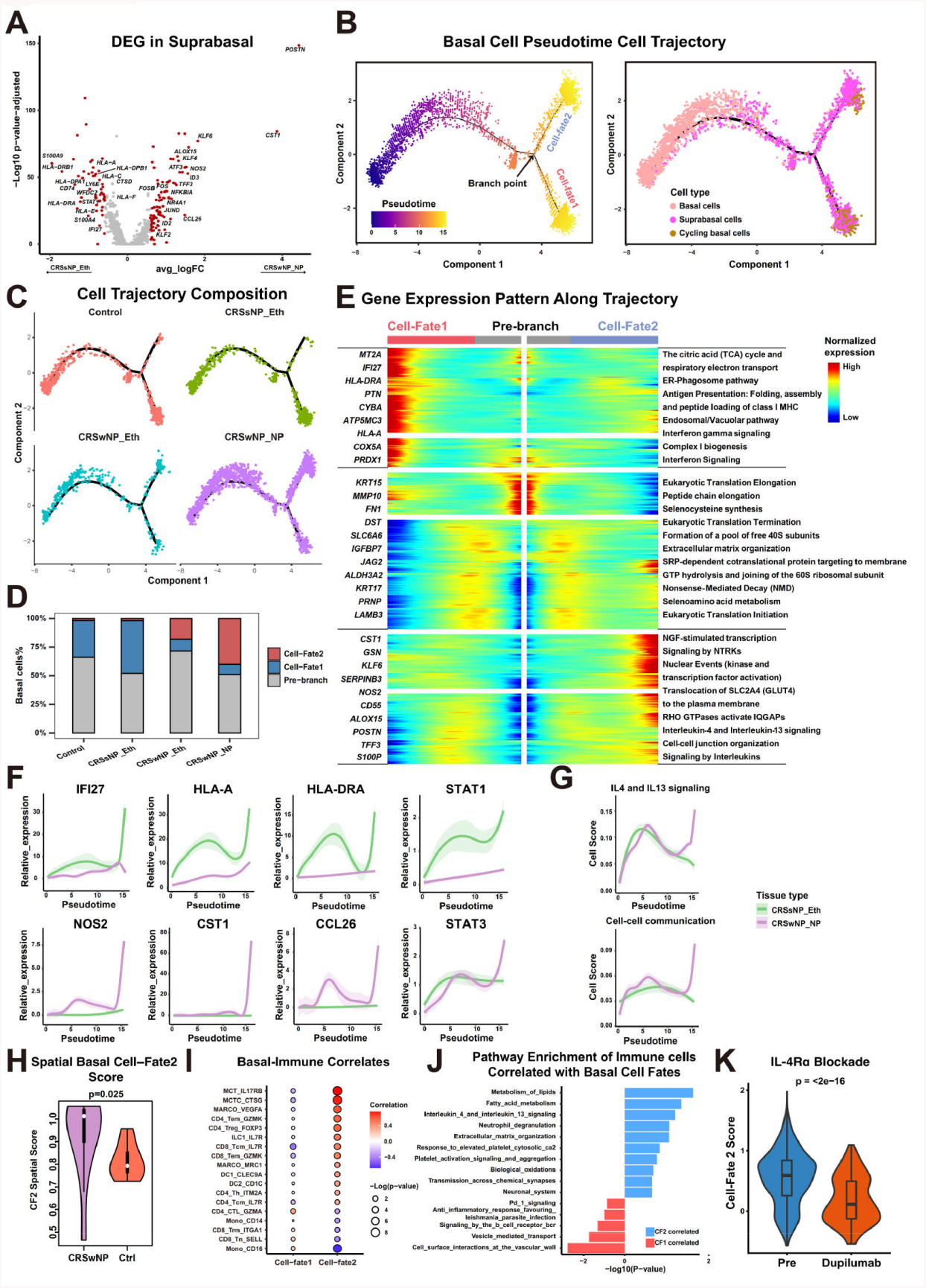
Nascent Basal Cells in Nasal Polyps Exhibit a Unique Transition Trajectory and Induce T2 Immune Response. (A) Volcano plot depicting differentially expressed genes in suprabasal cells between CRS nasal polyps and CRS without nasal polyps. The most significant genes are highlighted in red (|*F oldchange*| > 1.5). (B) Pseudotime trajectory analysis for basal cells using Monocle (left panel), accompanied by a cell density plot of the three basal cell subtypes along the pseudotime axis (right panel). (C) Cell density plot illustrating the distribution of basal cells from CRS and control samples along the pseudotime trajectory. (D) Histogram displaying the distribution of basal cells from CRS and control samples in three phases identified in (B). (E) Gene expression dynamics along the basal cell trajectory outlined in (B), from the pre-branch phase to cell fate 1 and cell fate 2. Genes are clustered into three gene sets, each characterized by specific expression profiles, as demonstrated by marker genes (left) and enriched pathways (right) unique to each cluster. (F) Dynamic expression of genes upregulated in CRS nasal polyps (top panels) and CRS without nasal polyps (bottom panels) during basal cell transition along pseudotime in CRS nasal polyps (purple) and CRS without nasal polyps (green). (G) Dynamic expression score of functional pathway signatures upregulated in CRS nasal polyps during basal cell transition along pseudotime in CRS nasal polyps (purple) and CRS without nasal polyps (green). (H) Violin plots comparing expression scores of Cell-fate2 basal cell signature between CRS nasal polyps (purple) and healthy control samples (red) in DSP data within PanCK+ regions. (I) Dotplot illustrating the correlation between different cell-fate basal cells and immune cells. Correlations with a p-value < 0.2 are displayed. (J) Pathways enriched in the top 5 cells correlated with Cell-fate1/2 basal cells, based on GSEA analysis using the REACTOME gene set. (K) Violin plots comparing the expression scores of Cell-fate2 basal cell signature between basal cells in pre-treatment (blue) and post-treatment (red) nasal polyps samples from an individual treated with IL-4Rα antibody.

We stratified Cell-fate1 and Cell-fate2 to reveal important differences in genes and pathways associated with each basal cell fate (Fig.6E-F), including an enrichment of IL4 and IL13 signaling, and cell-cell communication in CRSwNP (Fig.6G), in contrast to heightened IFN signaling and antigen presentation seen in CRSsNP (Fig.S6C). Cell-fate2 for basal cells also correlated with multiple metabolic, immune attractant, and tissue remodeling pathways (Fig.6E). A potent link between Cell-fate2 basal cells and eosinophil infiltration was further delineated by spatial transcriptomic analysis (Fig.S6D). Spatial transcriptomic (Fig.6H) and reconstruction of the pseudotime tracjectory also confirmed the enrichment of Cell-fate2 basal cells in CRSwNP tissues (Fig.S6E-F), and further highlighted the deviation towards key basal Cell-fate2 pathways in CRSwNP(Fig.S6G-H). We observed an increase in basal-immune cell interactions from scRNA-seq in Cell-fate2 directed basal cells (Fig.6I), and increased enrichment of pathways related to metabolism, IL4/IL13 signaling, neutrophil degranulation, and tissue remodeling (Fig.6J). These results suggest that basal cells from CRSwNP patients may differentiate towards a cellular state that is more conducive for immune system co-mingling along with tissue remodeling such as polyp formation, implicating basal cells and this Cell-fate2 differentiation pathway as a pivotal determinant for NP formation through epithelial-immune signaling and remodeling.

### A Reduction in the Cell-Fate2 Basal Cell Trajectory Upon Use of Immunotherapeutics Intervention for CRSwNP

The upregulation of *IL4* and *IL13* in CRSwNP disease, and in basal Cell-fate2 trajectory, implicates the central role of basal cells in coordinating CRSwNP and NP development. This was further supported by results from IL4 and IL13 cytokine stimulation of non-NP derived basal cells (10), indicating a skew towards the Cell-fate2 signature (Fig.S6I). Dupilumab is an IL-4/-13 receptor alpha antagonist that is FDA-approved as a primary and/or maintenance treatment in adult patients with poorly controlled CRSwNP (23, 24). Inferior turbinate and NP tissues sampled pre- and post-dupilumab treatment were reanalyzed using scRNA-seq (10), and found to have a statistically significant reduction in Cell-fate2 transcriptomic signature in basal cells (Fig.6K). Taken together, these results clarify the role of basal cells and the Cell-fate2 developmental trajectory as the center of both epithelial-immune system interactions and remodeling in NP formation in patients suffering from chronic rhinosinusitis.

## Discussion

The present study provides an in-depth analysis of the complex immune and epithelial landscape in chronic rhinosinusitis (CRS) without and without nasal polyps, through singlecell transcriptomic profiling, and orthogonal interrogation of the intact tissue microenvironment with spatial transcriptomics. Our findings begin to unravel intricate immune-epithelial interactions and remodeling at play in nasal polyp tissues, thus shedding light on the cellular and molecular mechanisms that drive the pathogenesis of CRS, particularly related to NP formation.

In CRSwNP disease, our data outlined a role for macrophage polarization and recruitment of eosinophils into the epithelial compartment (Fig.2), Type II inflammatory activation in T cells (Fig.3), *IL4* and *IL13* activation in MCs and interactions with CD4+ T cells (Fig.4), an epithelial-immune axis harbored by Tuft cells (Fig.5) and basal cells (Fig.6), and a unique differential pathway for basal cells associated with NP formation (Fig.6). Notably, we observed polarization of macrophages towards an M2 phenotype specifically in CRSwNP that primes Type 2 inflammation. The M2 macrophages were found to secrete *CCL13* and *CCL18*, which are potent chemokines that promote eosinophilic infiltration into the upper airway epithelium. This observation emphasizes the role of macrophages in coordinating and molding the inflammatory milieu in inflammatory CRS nasal polyp disease, and their potential as a therapeutic target for modulating Type II inflammation.

These data also revealed the predominance of ‘immunosuppressive’ Type II-skewed CD4+ and CD8+ T cells within nasal polyps, further highlighting the crosstalk between macrophages and T cells in this common form of chronic sinonasal immunity. This interplay between immune cells within upper airway microenvironment suggests the presence of an intricate balance between pro-inflammatory and regulatory T cell subsets in distinct CRS fates, with potential implications for the development of immunomodulatory therapies targeting specific T cell subsets.

Another revealing finding was the enrichment of MCs within nasal polyp tissues, which strongly correlate with type II immune responses. We demonstrated that IL4/IL13-related ligand-receptor interactions between MCs and CD4+ T cells played a critical role in promoting Type II immunity in CRSwNP. This finding underlines the dance between innate MCs in mediating acquired T cell immune responses observed in chronic upper airway inflammation, and suggests that targeting MCs or their interactions with CD4+ T cells also represent a promising therapeutic strategy for possibly modulating type II immune responses in CRSwNP.

Our analysis further suggested a critical correlation between Tuft epithelial cells and Th2 lymphoid cells in nasal polyposis. Tuft cells were found to be involved in prostaglandin synthesis and regulation, with ALOX5 and PTGS1 expression mediating interactions between Tuft cells and CD4+ T cells that expressed the PTGDR2 receptor in CRSwNP. This immune-epithelial interaction suggests that targeting Tuft cells or their mediators could represent an novel avenue for blunting and/or modulating Th2 cell-driven inflammation in CRSwNP.

Finally, we demonstrated that nascent basal cells in nasal polyps exhibited a unique transitional trajectory that may induce Type II immune responses. The distinct Cell-fate2 basal cell trajectory identified within CRSwNP may provide a roadmap as to the aberrant epithelial regeneration observed in the mucosal tissues of these patients, with potential implications for understanding the tissue remodeling and immune-trafficking processes observed in CRSwNP, including that of NP generation. Experimental validation using IL4 and IL13 stimulation, or from dupilumab biologic treatment of a CRSwNP patient, further underscored the potential for targeting basal cell dynamics and the discrete interactions between epithelial progenitor cells and immunocyte populations as a novel treatment avenue.

These findings together serve to provide key insights into the epithelial-immune interactions within the tissue microenvironment of CRS, and their roles in tissue remodeling, immune cell attraction, and ultimately, NP formation in CRSwNP patients. By dissecting the subtle autocrine and paracrine cellular and molecular signaling interplay in CRS using higher-resolution tools, these multi-dimensional analyses implicate an array of pivotal actors and promising therapeutic targets for the modulation of both upper airway inflammation and tissue remodeling processes in chronic rhinosinusitis. Further research is needed to validate these findings in larger cohorts, and to explore the true therapeutic potential of decoupling immune-epithelial interactions in CRS. The multi-scaled transcriptomic resources generated herein will likely impact these future endeavors, and beyond.

## Materials & Methods

### Patient recruitment

Patients were diagnosed with CRSwNP and CRSsNP based on a European position paper on rhinosinusitis and nasal polyps (EPOS) 2012 and International Consensus of Allergy and Rhinology: Rhinosinusitis (ICAR:RS) guidelines. CRSwNP, CRSsNP, and controls were all recruited from Stanford University. Tissues from the ethmoid sinus mucosa or nasal polyps were collected during endoscopic sinus surgery. Five control patients underwent skull base surgery requiring ethmoid sinus surgery for treatment of cerebrospinal fluid leak, meningioma, or pituitary adenoma. None of the control patients had evidence of CRS or other upper airway inflammatory diseases on CT/MRI radiography or endoscopy. Patients with unilateral sinus disease, fungal or allergic fungal rhinosinusitis, antrochoanal polyps, cystic fibrosis, aspirin-exacerbated respiratory disease, or paranasal sinus cysts were excluded from this study. Patient characteristics, including demographics, medical history, and past medication use were collected. Patient data, including medication history, were independently verified through direct interview by a trained research technician/physician and by a questionnaire additionally administered on the day of surgery to confirm accuracy of existing records derived from patients’ electronic medical or pharmacy. In particular, to avoid confounders in the epithelial/immune cell findings associated with use of common anti-inflammatory medications in CRS, all included CRSsNP and CRSwNP patients were devoid of oral prednisone/methyl-prednisolone exposure and higher dose topical budesonide and mometasone nasal irrigations × 4 weeks, as well as lower-dose topical nasal steroid sprays such as fluticasone and mometasone for 2 weeks, prior to ethmoid or NP tissue sampling. Antibiotic use within 4 weeks of surgery also led to exclusion. Any doubt in patient medication use led to exclusion from final analysis. Patients’ characteristics are shown in Table 1. The study complied with the Declaration of Helsinki and all relevant ethical regulations of each institution, and written informed consent was obtained from each patient approved Institutional Review Board (IRB) protocols in accordance with the regulations of the Research Compliance Office at Stanford University (IRB 18981).

### Single-cell RNA sequencing and data processing

Each sample was received directly from surgeons and promptly delivered to the laboratory on ice. Upon arrival at the laboratory, the samples were immediately processed. The ethmoid sinus mucosa was removed from the bone and nasal polyps were left intact and were minced into small pieces by scissors on ice. The minced tissues were placed into a C tube (Miltenyi Biotec, Bergisch Gladbach, Germany) within a solution of RPMI 1640 (Gibco, Grand Island, NY) containing 10% fetal bovine serum (FBS), 0.02 mg/ml DNase I (Millipore Sigma, St. Louis, MO), and 4 mg/ml collagenase type IV (Thermo Fisher Scientific). The mixture was homogenized using the gentleMACS Dissociator (Miltenyi Biotec) and incubated at 37°C for total of 30 minutes (15 minutes, 2 times) with rotated using MACSmix Tube Rotator (Miltenyi Biotec). Between and after the two incubations, they were also homogenized in a gentleMACS Dissociator. Finally, the samples were filtered through a 70-µm cell strainer and spun down at 500g for 5 min. Red blood cells (RBC) were lysed using the RBC Lysis Solution (BioLegend, San Diego, CA) for 4 min at room temperature. Cells were then washed with ice-cold PBS and spun down at 500g for 5 min at 4°C before resuspension in RPMI containing 10

The single cell suspension was loaded onto the Chromium Controller (10x Genomics) using the Chromium single cell 3’ Reagent Kit v3 (10X Genomics), and scRNA-seq libraries generated in accordance with the manufacturer’s protocols. Sequencing was performed on a Illumina HiSeq 4000 with 75 bp pair end reads.

The CellRanger v3.1.0 (10X Genomics) analysis pipeline was used to generate a final digital expression matrix. Raw sequence reads were preprocessed and mapped onto the reference human genome (GRCh38-3.0.0). These data were used as input into the Seurat package (4.1.1) (https://github.com/satijalab/seurat) for further analyses in R (4.2.0). As part of the quality control metrics, genes detected (UMI count > 0) in less than three cells, and cells containing a small number of genes detected (UMI count < 200) or high mitochondrial genome transcript ratio (25%) were removed. After normalizing and identifying variable features for each sample independently, the data from all the patients were combined using the top 30 dimensions in ‘FindIntegrationAnchors()’ and ‘IntegrateData()’ functions.

### Unsupervised clustering and cell type identification

The normalized expression level was calculated for each gene by dividing the read counts for each cell by the total counts and multiplied by a scale factor of 1,000,000. The natural-log transformed values were taken as the final measurement of expression level for each gene in a specific cell. Based on the normalized expression level, we next selected a subset of genes that with high cell-to-cell variation in the dataset. Then, the principal component analysis (PCA) was performed on these variable genes. Following the results of PCA, Harmony was performed to correct the batch effect among samples (25), then an adequate number (30-40) determined by Elbowplot of principal components (PCs) were selected for dimensionality reduction and clustering. The UMap algorithm with a resolution parameter in a range of 0.1-0.8 was applied for dimensionality reduction and visualization (26). To identify marker genes that define a cluster, differential expression analysis was performed by comparing each single cluster to all other cells. To accelerate the computational time of differential expression analysis, genes with > 0.25 log-fold difference on average between the two groups of cells and detectable in more than 25% of cells in either of the two groups of cells were retained. Using the above differentially expressed genes, cells were annotated to different cell types according to their well-known canonical markers. All the above analysis was performed using the Seurat R package (v 4.1.1)(27)

### Differentially expressed genes analysis in scRNA-seq data

To define genes that may function in between CRS with and without nasal polyps, differential expression analysis in specific cell groups was performed using the ‘FindMarkers’ function implemented in the Seurat package. The Wilcoxon rank sum test with log-scaled fold change > 0.25 and adjusted P value < 0.05 (bonferroni correction) was performed to select differentially expressed genes.

### Pathway analysis

To reveal the potential biological functions of T cells in two types of CRS, GSEA was performed with R package ‘clusterProfiler’ and ‘ReactomePA’ to identify pathway enriched under the REACTOME gene sets released by MsigDB (28–31). In Tuft cells, differentially expressed genes identified between CRS with and without nasal polyps were used to perform WikiPathway enrichment (32). Pathways that have a BH-adjusted P value () smaller than were defined as being significantly enriched, and GSEA was performed to further validate the pathway enrichment.

### Definition and calculation of gene signature scores

To assess the functional status of speific cells, relative signatures were collected from published literature as follows. A M2 signature was used to define the functional phenotype of macrophages. An inflammatory signature (32), Th1 and Th2 signature (33, 34) were used to assess T cell functions. In scRNA data, expression scores of specific signatures were calculated using AddModuleScore in the Seurat package. To validate the interaction between basal cells and T2 immune response, the expression score and enrichment of cell fate signatures were accessed in public single cell and bulk RNA-seq datasets (10). All genes associated with each pathway score are available in Supp Table 2. Violinplot was adopted to present the scoring difference among different types of CRS and healthy control samples, and Wilcoxon rank-sum test was performed to indicate the statistical significance.

### Construction of cell developmental trajectory

The developmental trajectory of the basal cells was inferred using the Monocle2 package (10). The 10x Genomics sequencing data was first imported into Monocle2 in CellDataSet class, and the negative binomial distribution was chosen to model the reads count data. Differentially expressed genes across different cell populations were identified and selected as input features to construct the trajectory. Then, a Reversed Graph Embedding algorithm was performed to reduce the data’s dimensionality. With the expression data projected into a lower dimensional space, cells were ordered in pseudotime and trajectory was built to describe how cells transit from one state into another. After the cell trajectories were constructed, differentially expressed genes along the pseudotime trajectory separated by the branch point were detected using the ‘differentialGeneTest’ function. For each interested gene, the expression trend along the pseudotime was estimated using non-linear regression, and plotted with a curve chart.

### Inference of cell-cell communications

R package Cellchat (v1.5.0) was adopted to identify significant ligand-receptor pairs within different types of CRS samples (35). Ligand-receptor communication probabilities/strengths were computed, tested, compared and visualized on the samples of CRS with and without nasal polyps. The minimum communication cells threshold was set to 10 and other parameters were left as default.

### GeoMx-Digital Spatial Profiling

Samples collected for NanoString GeoMx-Digital Spatial Profiling were fixed in 10 Slides were deparaffinized and prepared according to the official NanoString GeoMx-NGS RNA Manual Slide Preparation protocol (36). In brief, slides were baked for 30 min at 60°C before washing in Xylene (3 × washes at 5 min each), 100% EtOH (2 × washes at 5 min each), 95% EtOH (1 × wash at 5 min) and in 1X PBS (1 × wash at 1 min). Slides then underwent heat induced epitope retrieval at 99°C for 10 min in Tris-EDTA retrieval buffer (eBioscience, 00-4956-58).

Slides were then digested by Protease K (0.1µg/ml) for 5 mins at 37°C, and then washed with 1X PBS. Subsequently, slides were fixed by 10% neutral buffered formalin (EMS Diasum, 15740-04) for 5 min at room temperature, then the fixation process was stopped by 5 mins of 1X NBF Stop Buffer wash, followed by 5 mins of 1X PBS wash. The NanoString DSP Human CTA detection probe cocktail was then applied to the slides and incubated overnight (18 hrs) at 37°C. After hybridization, slides were washed in Stringent Wash Buffer (2X SSC, 50% Formamide) 2 times, every 5 mins. Slides were then washed by 2X SSC twice, 2 mins each. Buffer W was then applied to the slides for 30 mins, followed by antibody staining for 1hrCD45 D9M8I, Cell Signaling Technologies), PanCK (AE1+AE3, Novus). Slides were then washed by 2X SSC twice, 5mins each, and stained with 500nM SYTO 13 for 15 min, then loaded on the GeoMx machine. For GeoMx DSP sample collection, we followed the instructions described in the GeoMx DSP instrument user manual (MAN-10088-03). Briefly, individual ROIs were then selected the areas immune cells aggregate and epithelium presented on the apical side of the tissues which includes ROI based on CD45 positive or PanCK positive masks were selected with the consent of two or more investigators. On average, the ROI sizes are approximately 45217 um2 for CD45+ regions and 37501 um2 for PanCK+ regions. After sample collection, the NanoString NGS library preparation kit was used: Each ROI was uniquely indexed using Illumina’s i5 × i7 dual-indexing system. In total, 4 µL of collected sample was used in a PCR reaction with 1 µM of i5 primer, 1 µM i7 primer, and 1 × NanoString library prep PCR Master Mix. PCR reaction conditions were 37°C for 30 min, 50 °C for 10 min, 95°C for 3 min, 18 cycles of 95°C for 15 s, 65 °C for 60 s, 68°C for 30s, and final extension of 68°C for 5min. Then the product was purified with two rounds of AMPure XP beads (Beckman Coulter) at 1.2 × bead-to-sample ratio. Libraries were paired-end sequenced (2 × 75) on a NextSeq550.

### Digital Spatial Profiling Data Analysis

Probes from the NanoString CTA panel were mapped and counted using the NanoString GeoMx Data Analysis software pipeline (36), using the FASTQ output from NGS sequencing. Thereafter, the data underwent quality control and normalization steps with the ‘Geomx-Tools’ software from NanoString: First, ROI and probes that did not meet the default QC requirement were filtered out and not used in the subsequent analysis. Next, raw probe counts were transferred into a gene-level count matrix by calculating the geometric mean of probes for each gene. Normalization of gene counts were then performed, with the ‘Q3 norm’ method in ‘Geomx-Tools’. The Q3 normed gene counts were then used for all subsequent downstream analysis. Mean levels of spatial region-specific gene expression or mean levels of spatial expression scores of specific signatures, and also their correlations were adopted to validate corresponding results or hypotheses. Apart from published signatures, differential expressed genes identified in scRNA data were also applied to validate cell phenotype and function in the DSP data, and spatial region-specific expression scores were calculated with ssGSEA using the GSVA package (37). The Wilcoxon rank sum test was performed to calculate the significance of differences between samples.

### Statistical Analysis

All data analyses were conducted in R 4.2.0. Statistical significance was defined as a two-sided P value of less than 0.05. The comparison of cell fractions, expression levels of marker genes and gene signature scores among different types of CRS and control samples were performed using Wilcoxon rank sum test. The correlation analyses were performed using Spearman’s correlation test.

## ACKNOWLEDGEMENTS

The authors thank members of the Jiang and Nayak laboratories for insightful discussions. S.J. is supported by NIH DP2AI171139, R01AI149672, a Gilead’s Research Scholars Program in Hematologic Malignancies, and the Bill & Melinda Gates Foundation INV-002704. J.V.N. is supported by R01HL151677, 1U54CA260517, and U19 AI171421. This article reflects the views of the authors and should not be construed as representing the views or policies of the institutions that provided funding.

S.J. has received speaking honorariums from Cell Signaling Technology unrelated to this work. G.P.N., is co-founder of IonPath Inc and Akoya Biosciences, Inc., inventor on patent US9909167, and is a Scientific Advisory Board member for Akoya Biosciences, Inc. The other authors declare no competing interests.

## AUTHOR CONTRIBUTIONS

Conceptualization: S.J., I.T.L., T.N., J.V.N.

Methodology and Analysis: G.L., T.N., I.T.L., B.Z, J.V.N., S.J.

Novel Reagents and Tools: D.T.B., C.H.Y., J.B.O., D.Z., S.S.D., P.A.G., A.Y., D.K., K.P., M.T.C., M.L., Z.M.P., P.H.H., D.W., J.C., Q.M., Z.L., G.P.N., D.B.

Writing – Original Draft: S.J., J.V.N, G.L., I.T.L., T.N., B.Z.

Writing – Reviewing and Editing: all authors

Supervision: S.J., J.V.N.

## Supplementary Figures

**Figure S1.**
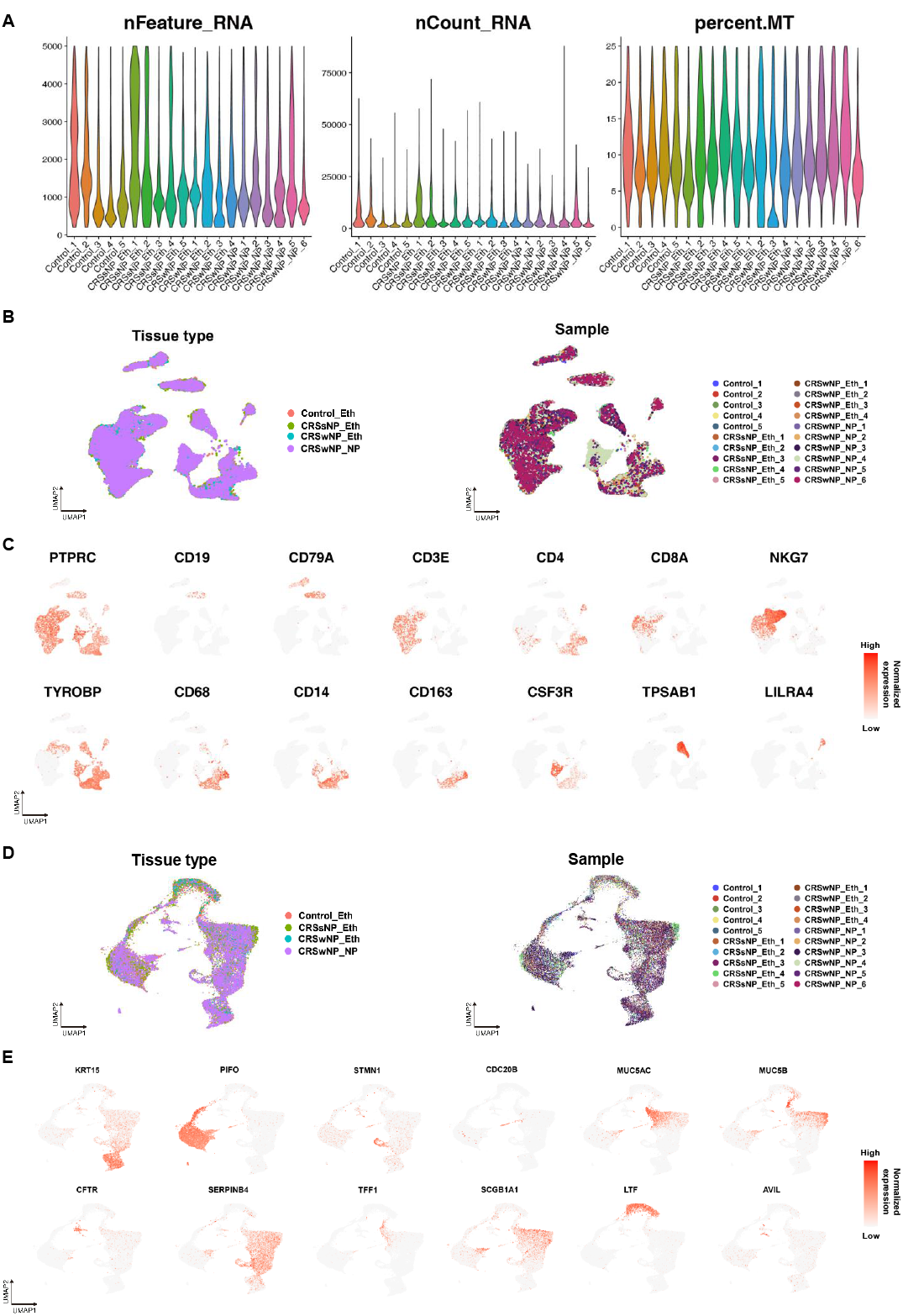
Comprehensive Single-Cell Transcriptomic Analysis Reveals the Complex Immune and Epithelial Microenvironment in CRS, related to Figure1. (A) Violin plots showing number of unique genes (left), number of total molecules (middle) and percentage of mitochondrial counts (right) of each cell in the single cell dataset. (B) UMAP plots showing immune cell origins by color, the origin of tissue types (left panel) and the origin of patient samples (right panel). (C) UMAP plot showing the expression of selected marker genes for the defined immune cell groups. (D) UMAP plots showing epithelial cell origins by color, the origin of tissue types (left panel) and the origin of patient samples (right panel). (E) UMAP plot showing the expression of selected marker genes for the defined epithelial cell groups.

**Figure S2.**
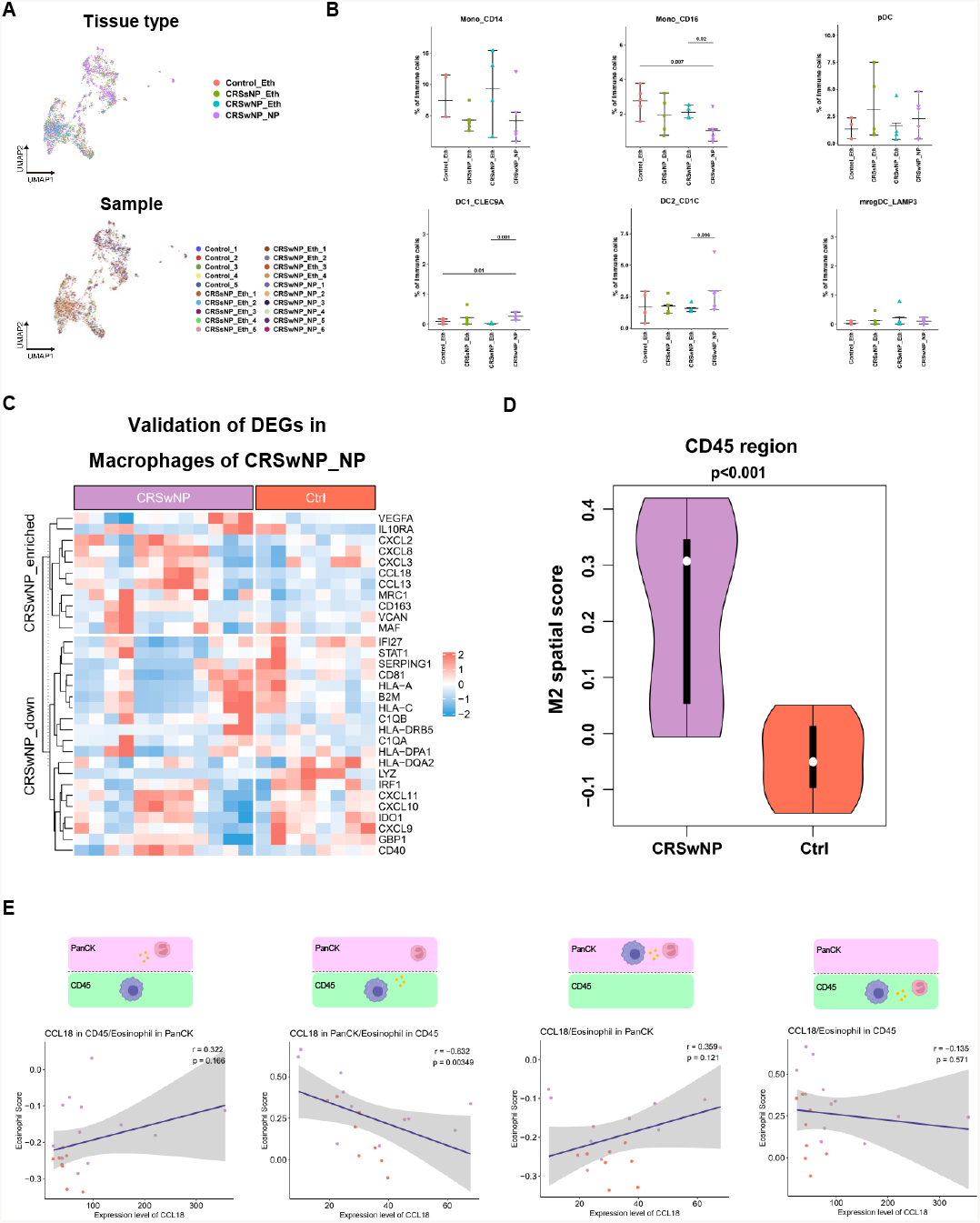
Polarization of Macrophages to M2 Phenotype Drives Type 2 Inflammation in CRS Nasal Polyps, related to Figure2. (A) UMAP plots showing myeloid cell origins by color, the origin of tissue types (upper panel) and the origin of patient samples (bottom panel). (B) Comparison of other myeloid cell fractions between CRS and control samples using the Wilcoxon test (two-sided). (C) Heatmap illustrating the normalized expression of genes upregulated and downregulated in nasal polyp macrophages in CD45+ regions of GeoMx data. (D) Violin plots comparing expression scores of M2 signature between CRS nasal polyps (purple) and healthy control samples (red) in GeoMx data within CD45+ regions. (E) Scatter plots illustrating the correlation between CCL18 expression levels and eosinophil signature scores in CD45+ or PanCK+ regions of GeoMx data, with sample origins color-coded to represent CRS nasal polyps (purple) and healthy control samples (orange). Diagrams above the scatter plots indicate regions where CCL18 and eosinophil spatial gene signatures were captured.

**Figure S3.**
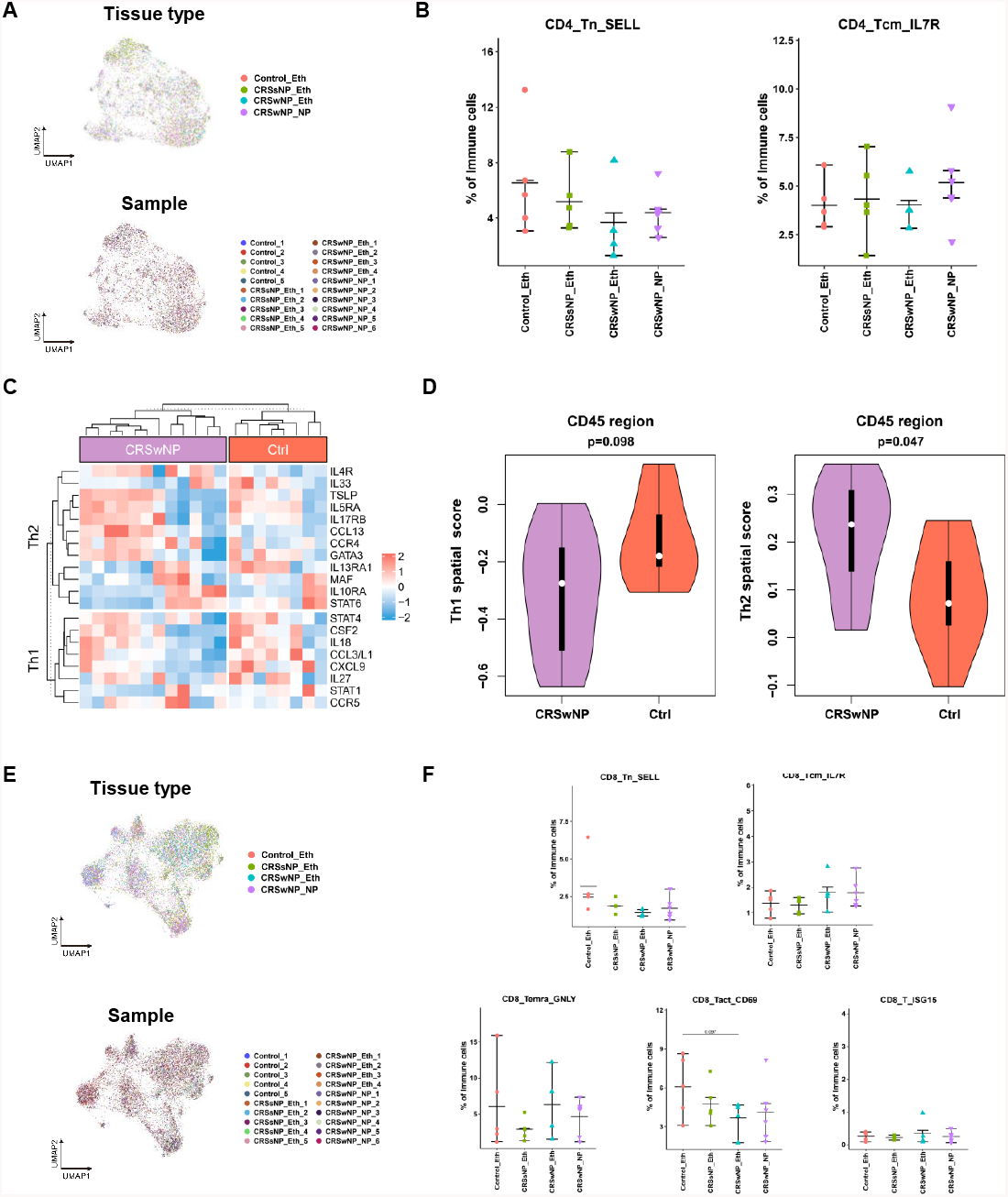
Regulatory CD4+ and CD8+ T Cells Predominate in Nasal Polyps, related to Figure3. (A) UMAP plots showing CD4+ T cell origins by color, the origin of tissue types (upper panel) and the origin of patient samples (bottom panel). (B) Comparison of other CD4+ T cell fractions between CRS and control samples using the Wilcoxon test (two-sided). (C) Heatmap illustrating the normalized expression of Th1/2 marker genes in CD45+ regions of GeoMx data. (D) Violin plots comparing expression scores of Th1/2 signatures between CRS nasal polyps (purple) and healthy control samples (red) in GeoMx data within CD45+ regions. (E) UMAP plots showing CD8+ T cell origins by color, the origin of tissue types (upper panel) and the origin of patient samples (bottom panel). (F) Comparison of other CD8+ T cell fractions between CRS and control samples using the Wilcoxon test (two-sided).

**Figure S4.**
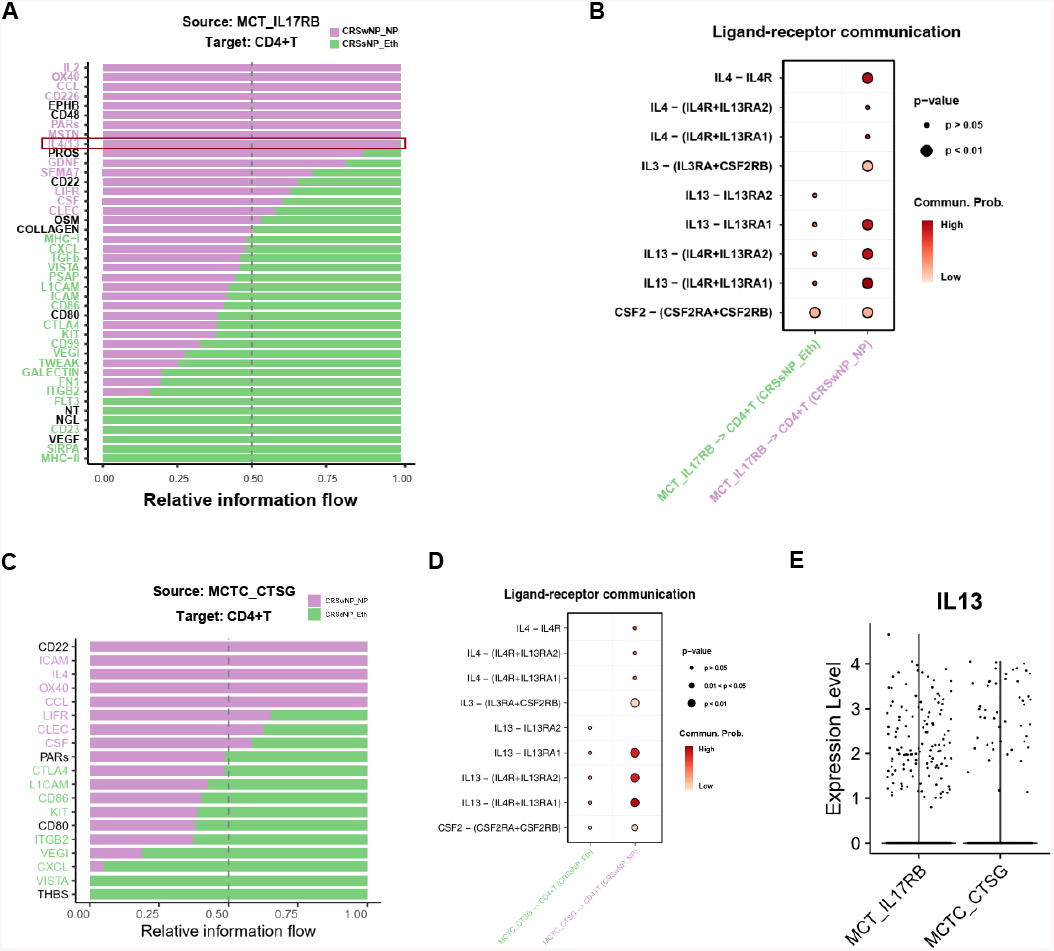
Mast Cell Enrichment in Nasal Polyps Correlates with Type 2 Immune Responses, related to Figure4. (A, C) Ligand-receptor (L-R) interactions identified between two subtypes of mast cells, MCT_IL17RB(A)/MCTC_CTSG(C) and CD4+ T cells in CRSwNP (purple) and CRSsNP (green). L-R pairs with purple bars crossing the 0.5 dotted line indicate predominance in CRSwNP, while those with green bars crossing the dotted line indicate predominance in CRSsNP. Significant interactions are color-coded accordingly (p<0.05, Wilcoxon test). (B, D) Dot plot demonstrating the significance and strength of IL4/IL13-related ligand-receptor interactions between two subtypes of mast cells, MCT_IL17RB(B)/MCTC_CTSG(D) and CD4+ T cells in CRSwNP (purple) and CRSsNP (green). (E) Scatter plots depicting IL4 and IL13 expression levels in mast cell subtypes, and their enrichment in MCT_IL17RB.

**Figure S5.**
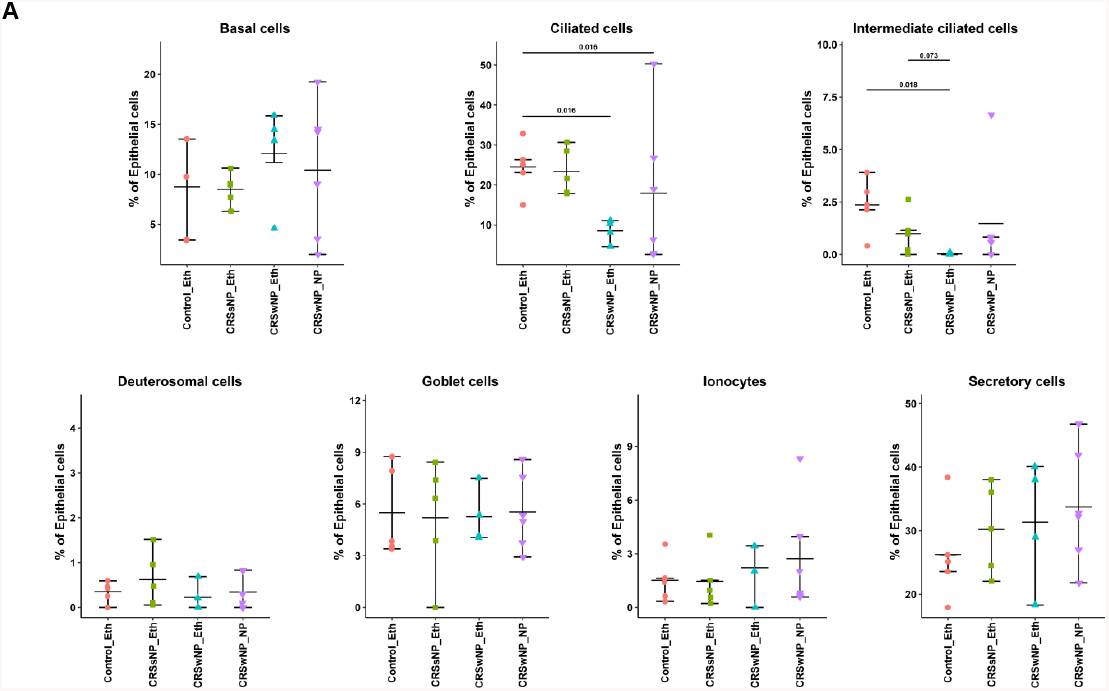
Epithelial composition difference in CRS, related to Figure5. Comparison of other epithelial cell fractions between CRS and control samples using the Wilcoxon test (two-sided).

**Figure S6.**
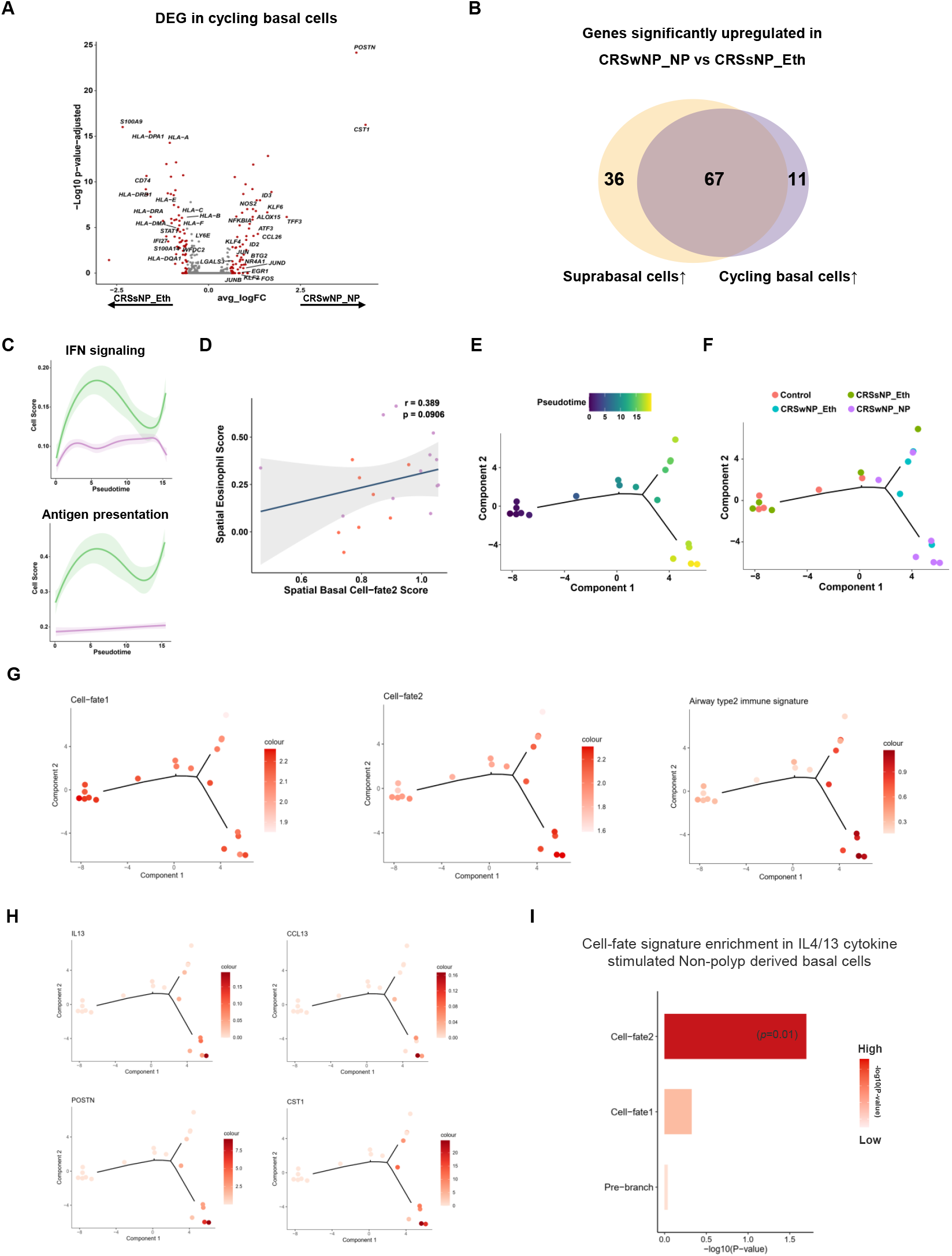
Nascent Basal Cells in Nasal Polyps Exhibit a Unique Transition Trajectory and Induce T2 Immune Response, related to Figure6. (A) Volcano plot depicting differentially expressed genes in cycling basal cells between CRS nasal polyps and CRS without nasal polyps. The most significant genes are highlighted in red (|Fold change| > 1.5). (B) Venn plot depicting overlap between upregulated genes in suprabasal cells and cycling cells in nasal polyps. (C) Dynamic expression score of functional pathway signatures upregulated in CRS without nasal polyps during basal cell transition along pseudotime in CRS nasal polyps (purple) and CRS without nasal polyps (green). (D) Scatter plot and regression line illustrating the correlation between Cell-fate2 basal cell spatial gene signature expression scores in PanCK+ regions and eosinophil cell spatial gene signature expression scores in CD45+ regions. Dots are colored to represent patient sample origins. The grey region indicates the confidence interval. The regression index and p-values are shown in the plots. (E) Pseudotime trajectory analysis for pseudo-bulk data of each sample in the single cell dataset using differentially expressed genes among the three branches in Figure 6B for ordering. (F) Cell density plot illustrating the distribution of CRS and control samples along the pseudotime trajectory. (G) Pseudotime plot showing the expression of basal cell-fate signatures for pseudo-bulk samples along the trajectory. (H) Pseudotime plot showing the expression of genes upregulated in CRS nasal polyps during basal cell transition along the trajectory. (I) Barplot showing the enrichment of basal cell-fate signature in IL4/13 cytokine stimulated Non-polyp derived basal cells compared with Non-polyp derived basal cell baseline.

